# Scanning the RBD-ACE2 molecular interactions in Omicron variant

**DOI:** 10.1101/2021.12.12.472253

**Authors:** Soumya Lipsa Rath, Aditya K. Padhi, Nabanita Mandal

## Abstract

The emergence of new SARS-CoV-2 variants poses a threat to the human population where it is difficult to assess the severity of a particular variant of the virus. Spike protein and specifically its receptor binding domain (RBD) which makes direct interaction with the ACE2 receptor of the human has shown prominent amino acid substitutions in most of the Variants of Concern. Here, by using all-atom molecular dynamics simulations we compare the interaction of Wild-type RBD/ACE2 receptor complex with that of the latest Omicron variant of the virus. We observed a very interesting diversification of the charge, dynamics and energetics of the protein complex formed upon mutations. These results would help us in understanding the molecular basis of binding of the Omicron variant with that of SARS-CoV-2 Wild-type.

**TOC Graphics:** 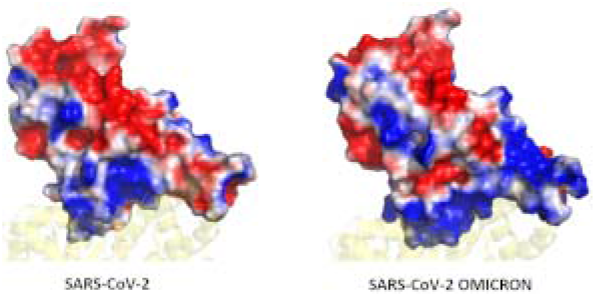

---

The Covid19 pandemic started in the later part of 2019 claiming millions of lives and affecting several others physically and mentally^1, 2^. Although several vaccines have been developed against the virus, we still have to remain observant due to emergence of novel variants of the virus. It is in the nature of RNA viruses, such as coronaviruses, to undergo frequent genomic mutations^3^. As a result of the mutation, the structure as well as the dynamics of the proteins, including the viral proteins get affected ^4, 5^. The WHO has briefly classified the variants of SARS-CoV-2 into Variants of Concern (VOC) and Variants of Interest (VOI). The VOCs are of particular interest due to higher infectivity and transmissibility^2^. After the Alpha, Beta, Gamma and Delta variants, recently, a new Omicron variant (B.1.1.529) has been identified^2^. The initial reports indicate a notable 32 mutations in the Spike glycoprotein of the virus^6^. The Spike glycoprotein is one of the largest structural proteins of the virus, whose stalk is embedded in the viral membrane while a large head interacts with the host Angiotensin Converting Enzyme 2 (ACE2) receptor^7^. The primary role of the Spike protein lies in the attachment of the virus to the host cells^8^.

The Spike protein, constituted by three peptide chains, has a specific domain on the surface which is known as the Receptor Binding Domain or RBD^9^. This RBD is the interacting domain of the virus. Most of the reported mutations of VOCs have been found to occur in the RBD which vary from residue 333-527^10^. For instance in N501Y mutate in Alpha, Beta and Gamma variants; K417T in Alpha and Beta; and T478K in the Delta variant^2^. As per the initial reports, a total of 15 mutations have been identified on the RBD of the Omicron variant. Since the primary role of RBD lies in the binding with the ACE2 receptor, the identified mutations would also influence the nature of interaction in the protein-protein complex as observed in earlier studies^11^. Molecular studies have shown that mutations have resulted in variation in the residue wise interaction energies reported for SARS and SARS-CoV-2 as well as in other variants^12,13^.

In our study we have considered the RBD/ACE2 complex taken from the crystal structure as the Wild-type system^14^. Due to the lack of available crystal structures of the Omicron variant and the urgency in understanding the molecular details of protein interactions, we have generated the Omicron variant model using homology modeling using Modeller^15^. We have omitted the glycans in our study, since earlier studies by Amaro et. al. confirmed that the RBD of the Spike protein had significantly less glycans compared to the rest of the Spike protein and did not directly participate in the ACE2 interaction^16^. Subsequently both the systems were subjected to minimization, equilibration and production run using Gromacs MD Simulation package^17^ and CHARMM36 force field^18^ for 100 ns (ran in triplicates) to the accuracy of the structural-dynamics parameters ascertain the accuracy of the structural-dynamics parameters (details in SI). Although the RBD is the interacting domain of the Spike protein, the exact site of interaction lies between residues 438-506 known as the Receptor Binding Motif (RBM) ^10^. Structurally, the RBD comprises a twisted beta sheet and a small alpha helix, while the RBM is made up of four loops and two small beta strands^9^.

During the 100 ns simulation run time, we observed stability of both the systems (Figure 1a). Surprisingly, the Omicron system was more stable than the Wild-type despite the mutations. Detailed RMS fluctuations of the Cα atoms of the residues shows some interesting observations. Compared with earlier studies with Alpha, Beta, Gamma and Delta^13^, the RMSF values of the Omicron variant over the simulation time were lower than the Wild-type (Figure 1b, c). This is noteworthy since earlier mutational studies had shown the RMSF values to be either similar or higher than the Wild-type system. This phenomenon was observed for both the ACE2 (Figure 1b) as well as the RBD (Figure 1b) part of the Omicron protein complex clearly indicating the formation of a more stable protein complex. Not only the RBM in the RBD but the whole protein appears very relaxed with the ACE2 partner protein.

**Figure 1.**
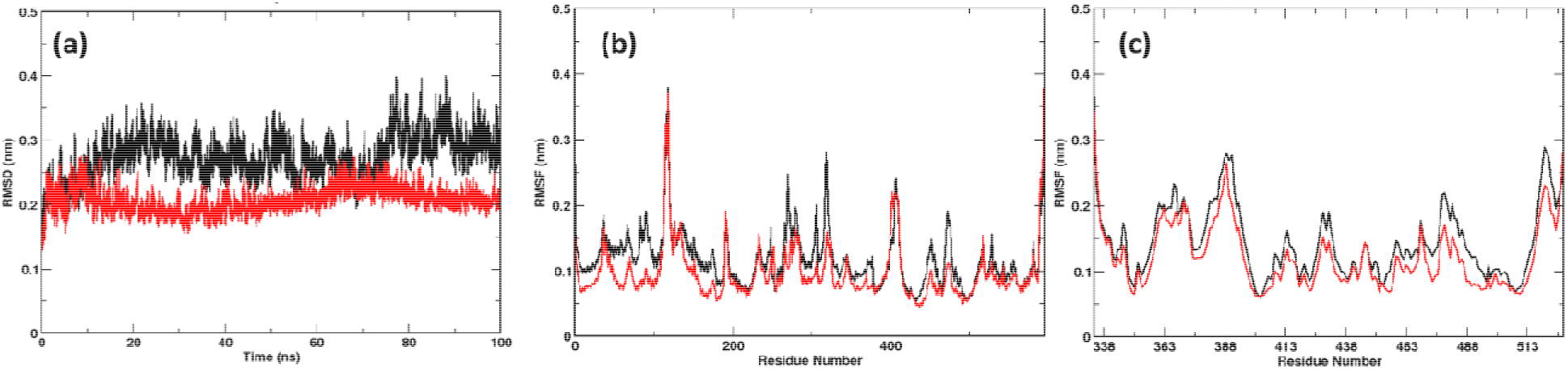
(a) RMSDs and RMSFs of (b) ACE2 and (c) RBD calculated over the simulation time. The Omicron system was found to be more stable than Wild-type. (Color Wild-type: black; Omicron: Red)

The peculiarity in the RMSF values, prompted us to investigate the dynamics of the RBD/ACE2 complex along the same lines. We performed Principal Component Analysis (PCA) of the Wild-type and Omicron RBD/ACE2 complex^19^. In Figure 2, a clear difference in the dynamics of the Omicron system was evident. In the Wild-type system the flexibility was found to be more along the first principal component. The trace values of 13.91 in Wild-type and 10.02 in Omicron also indicate compactness of the Omicron system.

**Figure 2.**
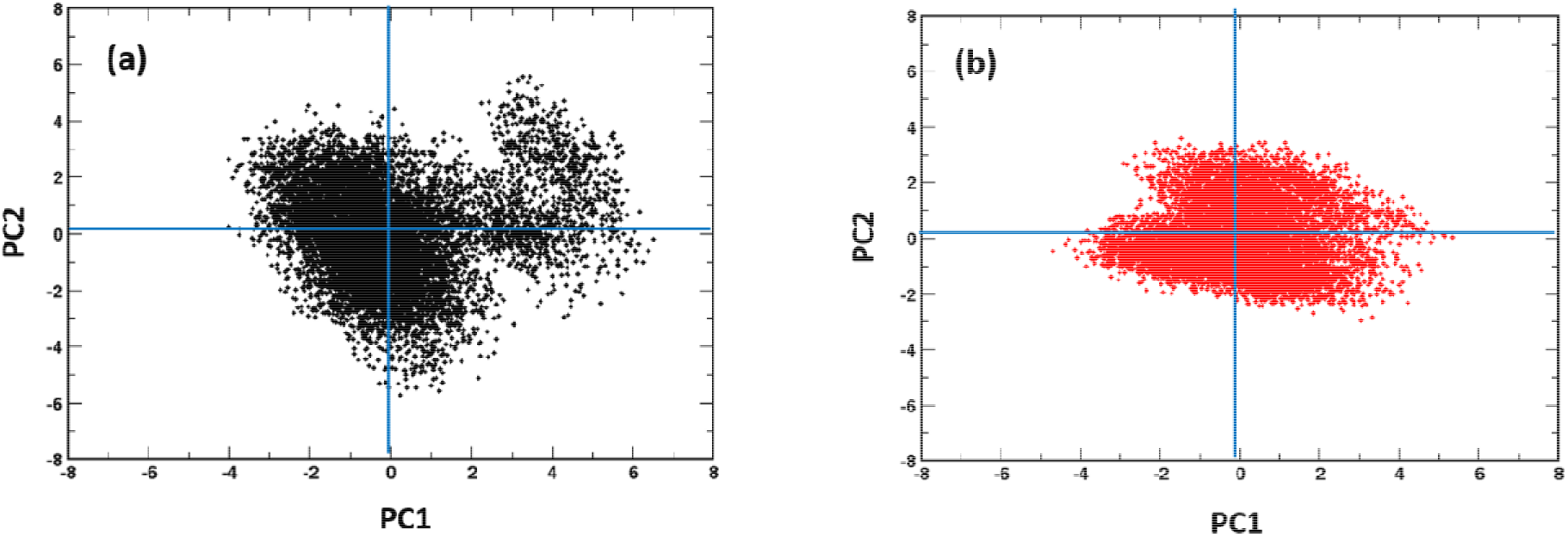
Projection of the first and second principal components of the Wild-type and Omicron system displaying the difference in RBD/ACE2 complex flexibility. (Color Wild-type: black; Omicron: Red)

The total interaction energies of the generated protein complex formed in both Wild-type RBD/ACE2 and Omicron RBD/ACE2 variants was compared during the last 10ns of the simulation time (Table 1). A value of −2658 kJ/mol of the Omicron distinctly shows the strength of the formed protein complex. Wild-type complex shows a value of only −1022 kJ/mol, which is 2.5 times lower than the Omicron system. Major changes were observed in the electrostatic energies of the protein. This can be explained due to the change in the amino acid R-groups in the mutating residues as shown in Table 2.

**Table 1.** Total energy of binding of RBD with the human ACE2 receptor

|  | Wild-type | Omicron |
| --- | --- | --- |
| van der Waal energy | -363.775 +/- 20.446 kJ/mol | -335.186 +/- 21.937 kJ/mol |
| Electrostatic energy | -1232.614 +/- 64.543 kJ/mol | -3083.962 +/- 89.676 kJ/mol |
| Polar solvation energy | 617.810 +/- 161.796 kJ/mol | 803.804 +/- 119.784 kJ/mol |
| SASA energy | -43.889 +/- 3.469 kJ/mol | -42.889 +/- 3.818 kJ/mol |
| Binding energy | -1022.467 +/- 150.703 kJ/mol | -2658.233 +/- 129.686 kJ/mol |

**Table 2.** Residue-wise binding energy calculated for the mutated residues in RBD

| Mutated Residue | Change in R-group | Wild-type (kJ/mol) | Omicron (kJ/mol) |
| --- | --- | --- | --- |
| ASP/GLY-339 | Negative to Neutral | 1.7602 | 157.4091 |
| LEU/SER-371 | Neutral to Positive | 2.6324 | 1.5017 |
| PRO/SER-373 | Neutral to Positive | 2.6935 | 2.7563 |
| PHE/SER-375 | Neutral to Positive | -0.2065 | -0.0565 |
| ASN/LYS-417 | Negative to Positive | -230.7012 | -4.1475 |
| LYS/ASN-440 | Positive to Negative | 2.7283 | -240.3853 |
| SER/GLY-446 | Positive to Neutral | -2.5231 | -1.1289 |
| ASN/SER-477 | Negative to Positive | -2.5193 | -0.3975 |
| LYS/THR-478 | Positive to Positive | 0.652 | -206.7846 |
| ALA/GLU-484 | Neutral to Negative | 233.7813 | -0.4936 |
| LYS/GLN-493 | Positive to Negative | -0.3732 | -222.7693 |
| SER/GLY-496 | Positive to Neutral | 1.2866 | 3.175 |
| ARG/GLN-498 | Positive to Negative | -0.2806 | -265.8465 |
| TYR/ASN-501 | Positive to Negative | -7.8806 | -14.9393 |
| HIS/TYR-505 | Positive to Positive | -16.9694 | -11.9467 |

The interface of RBD is lined by a number of charged residues such as ASP, GLU, LYS. Upon analyzing the mutations, we observe that several mutations resulted in change in the amino acid which could severely impact its binding potential. We calculated the per residue binding energies of Wild-type and Omicron variants using MM/PBSA. Since the RBD region alone shelters fifteen different mutations, we compared the binding energies of those mutated residues. There are three instances of mutations changing the residue from charged to neutral. While S496G and S446G do not show significant difference in the binding energies, the D339G mutation from a highly negative to a neutral residue makes it unfavorable for RBD to bind to the ACE2 (refer to Table S1). Similarly, not much difference could be seen for mutations that result in changing the residues from neutral to charged residues. Although, the change of A484E is favorable for the Omicron RBD to bind to ACE2. The most evident changes were seen when there is change of positive residues to negative such as K440N, K493Q and R498Q (Refer to Table S1). Here, more than 200kJ/mol gain in per residue binding energy could be seen. Conversely, when mutation led to change in amino acid from negative to positive there was a loss in binding energy values. A significant difference in binding energy was observed when highly positive K478 mutates to polar threonine residue which adds around −206 kJ/mol towards the binding of the RBD in the Omicron system. Apart from the mutated residues, we observed changes in binding energies of residues R454 and K458 in the RBM, where the Omicron system had a higher energy of binding (Table S1).

We also compared the difference in binding energies of the ACE2 residues (specifically the interface residues) and those that directly interact with the RBD. Those residues whose energies vary significantly are shown in Figure 3a. Here, we found reduced energies of residues S19, K26, K31, K68, K74 and K94. However, negatively charged residues E23, D30, E35, E37, D38, E56, E57, D67, E75, E87, all show an improved binding potential. The bias in the increase in binding energies of negatively charged residues instigated us to investigate the surface potential of the RBM in both the systems. Surface electrostatics of the RBM were calculated using the Adaptive Poisson Boltzmann Surface Area^20^. Figure 3b and c show a clear difference in the interfacial surface potential of the RBD in both the complexes. The Wild-type had shown both positively and negatively charged regions at the binding surface. Conversely, the Omicron variant had a highly positive binding interface. This corresponds very well with the increase in binding energies of the negatively charged residues present at the ACE2 interface.

**Figure 3.**
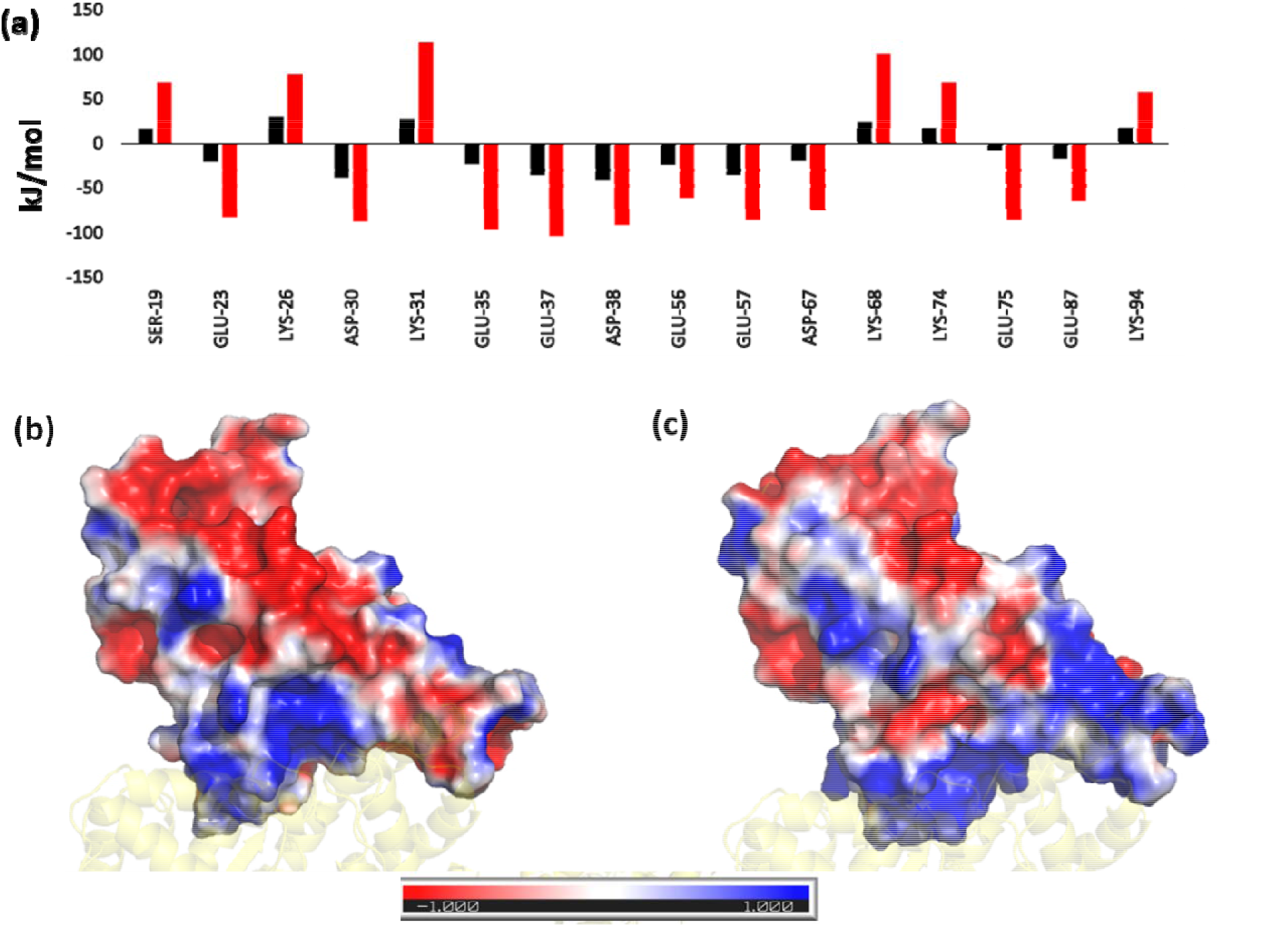
(a) Residue-wise binding energy of the ACE2 receptor highlighting those residues that show the most changes (Color coding similar to Figure 1). Electrostatic potential Map of the RBD in (a) Wild-type and (c) Omicron Variant displaying the difference in charge distribution. The red and blue colors denote the negative and positive potentials respectively.

Superimposition of the final conformations of both the complexes show an RMSD value of 1.44 Å, implying not much change in the protein conformation. The mode of binding was also found to be similar after running the simulation for 100ns. However, we observed a slight displacement of the N-terminal helix of ACE2 towards RBD in the Omicron complex. Projections of the principal modes using porcupine plots show that both RBD and ACE2 are highly dynamic in the Wild-type (Figure 4). Two different movements were observed along the first principal mode; the RBD and N-terminal ACE2 receptor of Wild-type move synchronously in clockwise direction. The second motion was of the C-terminal domain of ACE2 in the anticlockwise direction. The Omicron system, conversely shows remarkable reduction in the dynamics of Cα atoms (Figure 4b). The magnitude as well as the direction of the principal modes were different. The inter and intramolecular movements were primarily observed in the Omicron complex. The magnitude of clockwise movement of RBD and N-terminal portion of ACE2 was also found to have drastically reduced. This indicates the stability of the Omicron RBD/ACE2 complex similar to the Figure 2 above.

**Figure 4.**
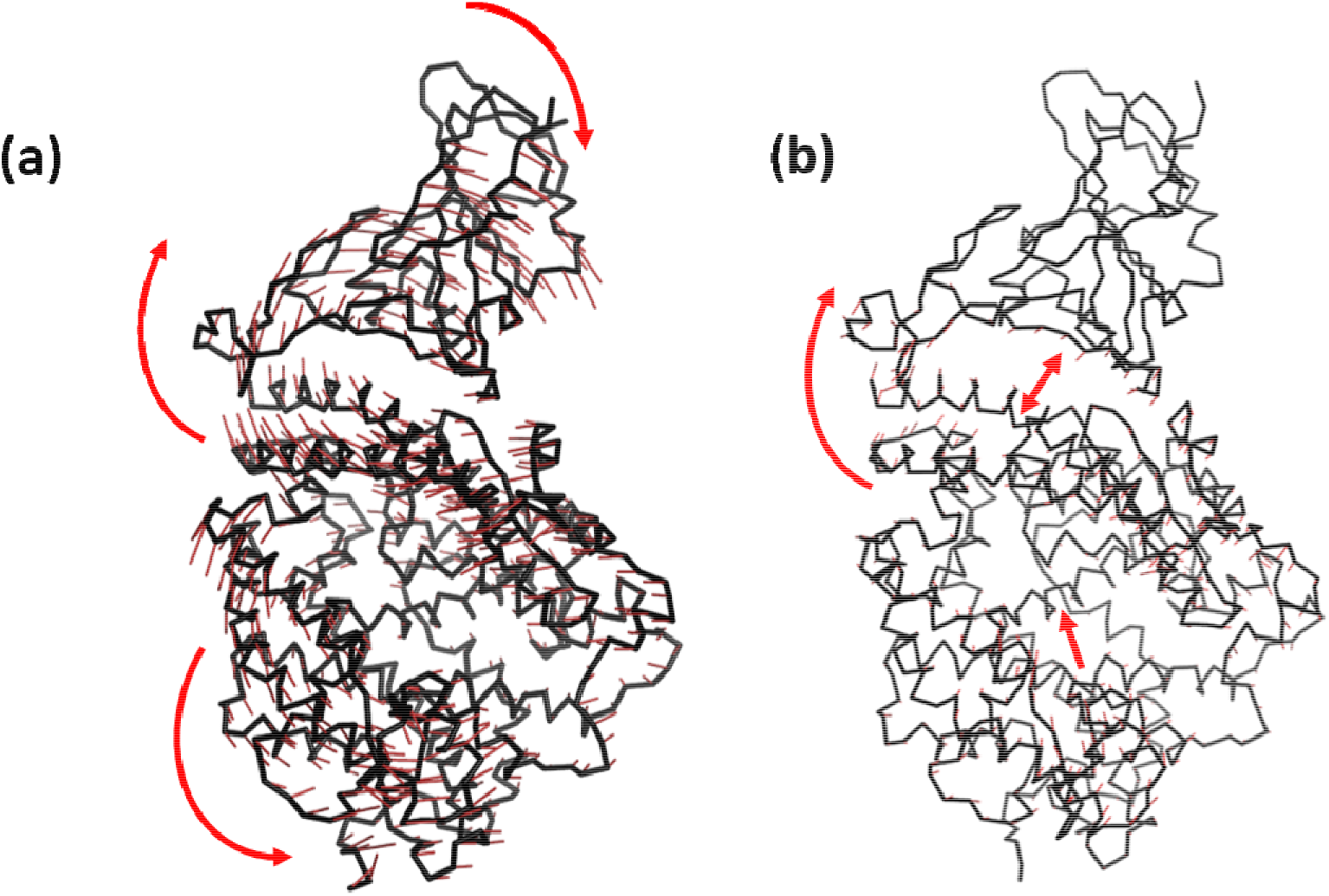
Porcupine plots constructed by projecting the displacement of the backbone CA atom along the first principal component in (a) Wild-type and (b) Omicron variant. The red arrow denote the direction of movements of the atoms

The relative orientation of the interfacial residues was also compared to gain more insights on the binding modes of both the proteins (Figure 5). We found that four of the mutated residues do not directly interact with the ACE2. Moreover, the mutated residues namely L371S, P373S and F375S do not show significant change in their energetic contributions. The D339S mutation was found to be highly repulsive, however since the residue does not directly interact with the ACE2, it doesn’t impact the protein complex formed. The most prominent changes were found for N417K, K440N, K478T, A484D, K493Q and R498Q. Except the N417K mutation which decreases the interaction energy, all the remaining residues massively increase the interaction energy of the RBD.

**Figure 5.**
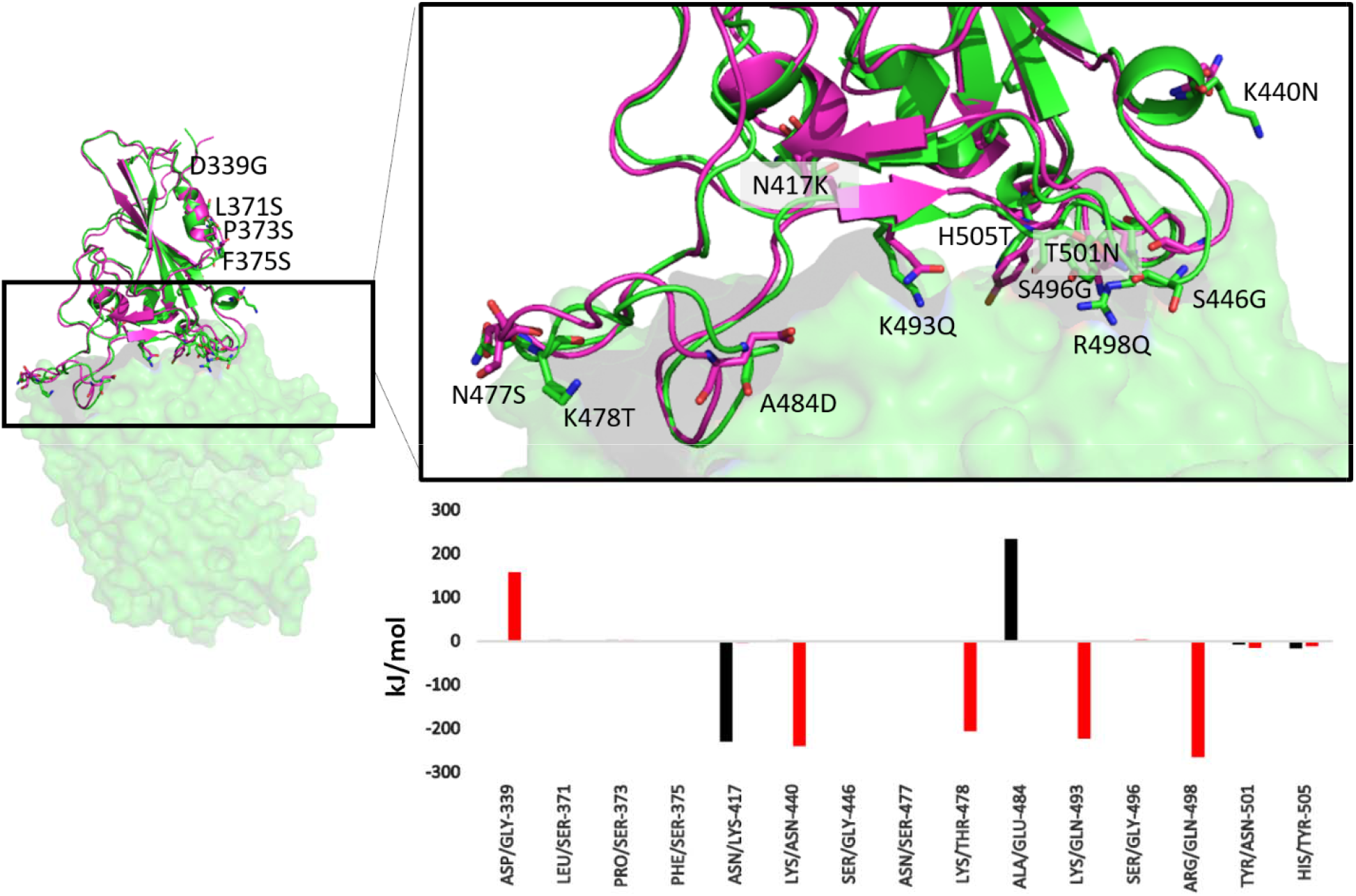
Superimposed structure of Wild-type (magenta) and Omicron (green) showing the mutated residues in stick representation and colored by CPK. The residue-wise binding energy is shown as a bar graph (Color scheme similar to Figure 1).

Earlier studies have emphasized the difference in dynamics of SARS-CoV and SARS-CoV-2 on the interfacial interactions between RBD and ACE2 [15]. To elucidate the difference in hydrogen bonding and hydrophobic interactions among both the systems we compared the interfacial interactions by using Ligplot+ ^21^ (Figure 6). Here we observed ten hydrogen bonded interactions in Wild-type while only eight in the Omicron variant (Figure 6a,b). Apart from a common H-bond between D483 of RBD and Y83 of ACE2, most of the H-bonds formed were remarkably different. The hydrophobic interactions were also observed to be relatively more in the Wild-type RBD/ACE2 complex. However, the number of residues participating in hydrophobic interactions in both RBD and ACE2 were higher in Omicron as can be seen in Figure 6c and d.

**Figure 6.**
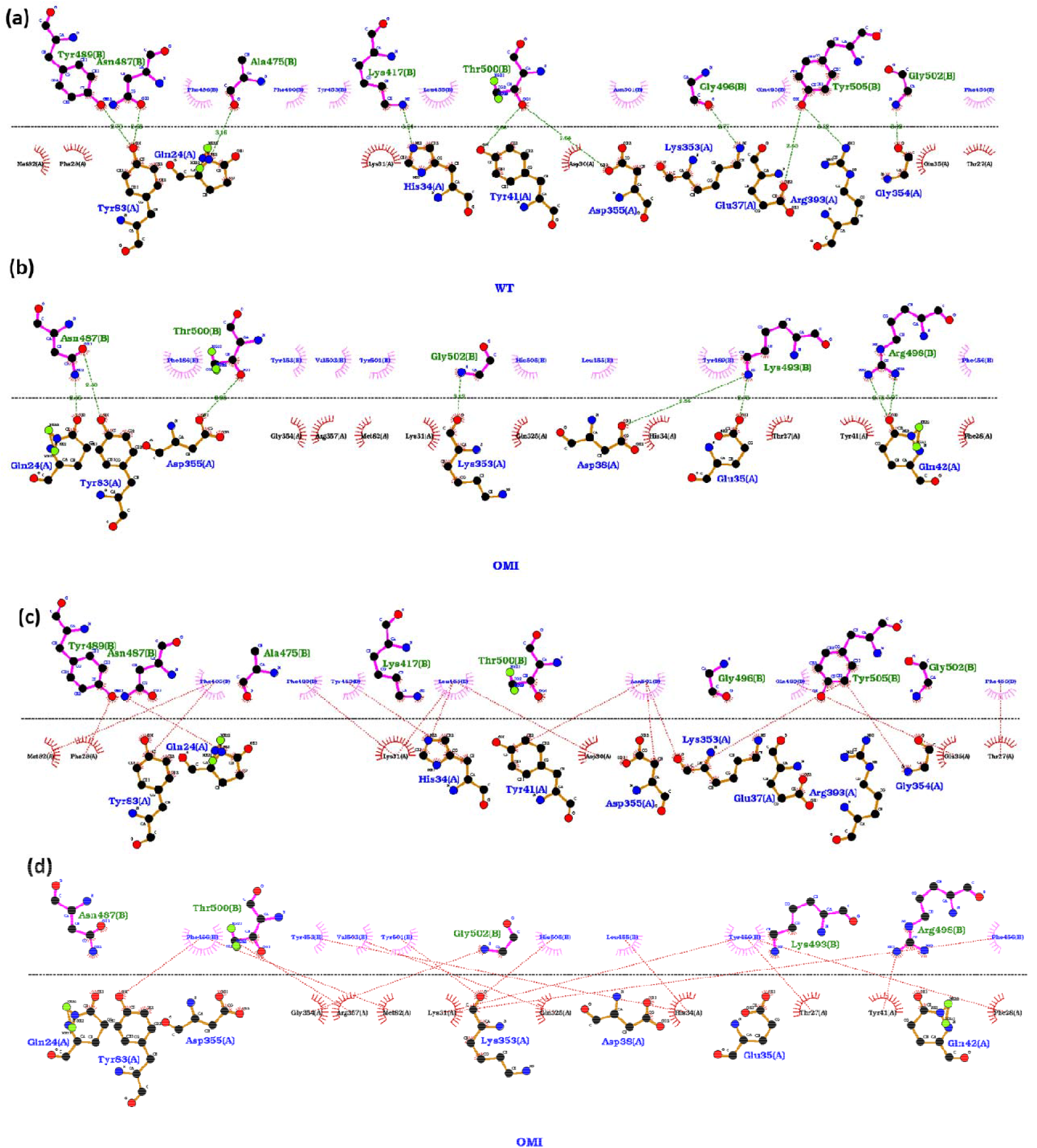
Interaction between RBD and ACE2 in (a) (c) Wild-type and (b) (d) Omicron variants. Hydrogen bonded interactions are shown in (a) and (b) and hydrophobic interactions in the (c) and (d) panels.

To summarize, we have made a detailed molecular analysis of the binding of the RBD domain of Wild-type SARS-CoV-2 and Omicron Variant with the human ACE2 receptor. Enhanced binding with the human receptor is one of the crucial factors in transmissibility of the virus. Apart from atomistic details related to the binding mode, we have meticulously analyzed the residue wise interaction energies of the mutated residues. The complementary changes were also observed in the human ACE2 receptor. Detailed surface electrostatics imply the change in the nature of the binding interface to a highly positive patch in RBD of Omicron. This resulting change is favorable for the binding of ACE2 which is lined by several negatively charged ASP and GLU residues. The overall relaxed dynamics of the protein complex further supports the stability of the Omicron when compared to Wild-type. The study not only provides a first-hand rationale for the high rate of transmission of the variant but would also prove crucial for the drug development studies as well as in the designing of antibodies.

## Acknowledgements

SLR thanks NIT Warangal for research seed grant (P1131) and National Energy Research Scientific Computing Center of the Ernest Orlando Lawrence Berkeley National Laboratory, a DOE Office of Science User Facility supported by the Office of Science of the U.S. Department of Energy under Contract No. DE-AC02-05CH11231 and the Extreme Science and Engineering Discovery Environment (XSEDE). We extend our gratitude to the Covid19 HPC Consortium for providing resources and helping researchers work for a noble cause.

## Associated Content

### Supporting Information Available

Includes description of Materials and Methods and Table S1, Figures S1-S2. This information is available free of charge *via* the internet

